# NASTRA: Accurate analysis of short tandem repeat markers by nanopore sequencing with repeat-structure-aware algorithm

**DOI:** 10.1101/2023.11.04.565630

**Authors:** Zilin Ren, Jiarong Zhang, Yixiang Zhang, Tingting Yang, Pingping Sun, Jiguo Xue, Xiaochen Bo, Bo Zhou, Jiangwei Yan, Ming Ni

## Abstract

Forensic short-tandem repeats (STR) genetic markers are multi-allelic and widely utilized for individual identification, kinship testing, and cell-line authentication. Nanopore sequencing, known for its portability, is emerging as a promising approach for STR typing, facilitating real-time and in-field testing. However, its efficacy is often hampered by sequencing noise. Previous methods rely on alignment-based genotyping, necessitating known alleles, which limits their applicability to unknown alleles. Here, we introduced NASTRA, an innovative allele reference-free tool for precise germline analysis of STR genetic markers. NASTRA incorporates a recursive algorithm to infer repeat structures of allele sequences using only known repeat motifs. Our tests, conducted on 80 individual samples and 8 DNA standards, have demonstrated NASTRA’s exceptional 100% accuracy in genotyping nearly all diploid STRs across various multiplex kits and flow cells. It surpasses alignment-based methods in accuracy and speed. In a paternity testing case study, NASTRA accurately identified three relationships among six individuals within an 18-minute sequencing duration. These results underscore NASTRA’s ability to perform STR analysis on both NGS and nanopore sequencing platforms, significantly enhancing the utility of nanopore sequencing in relevant applications.

**GRAPHICAL ABSTRACT:** 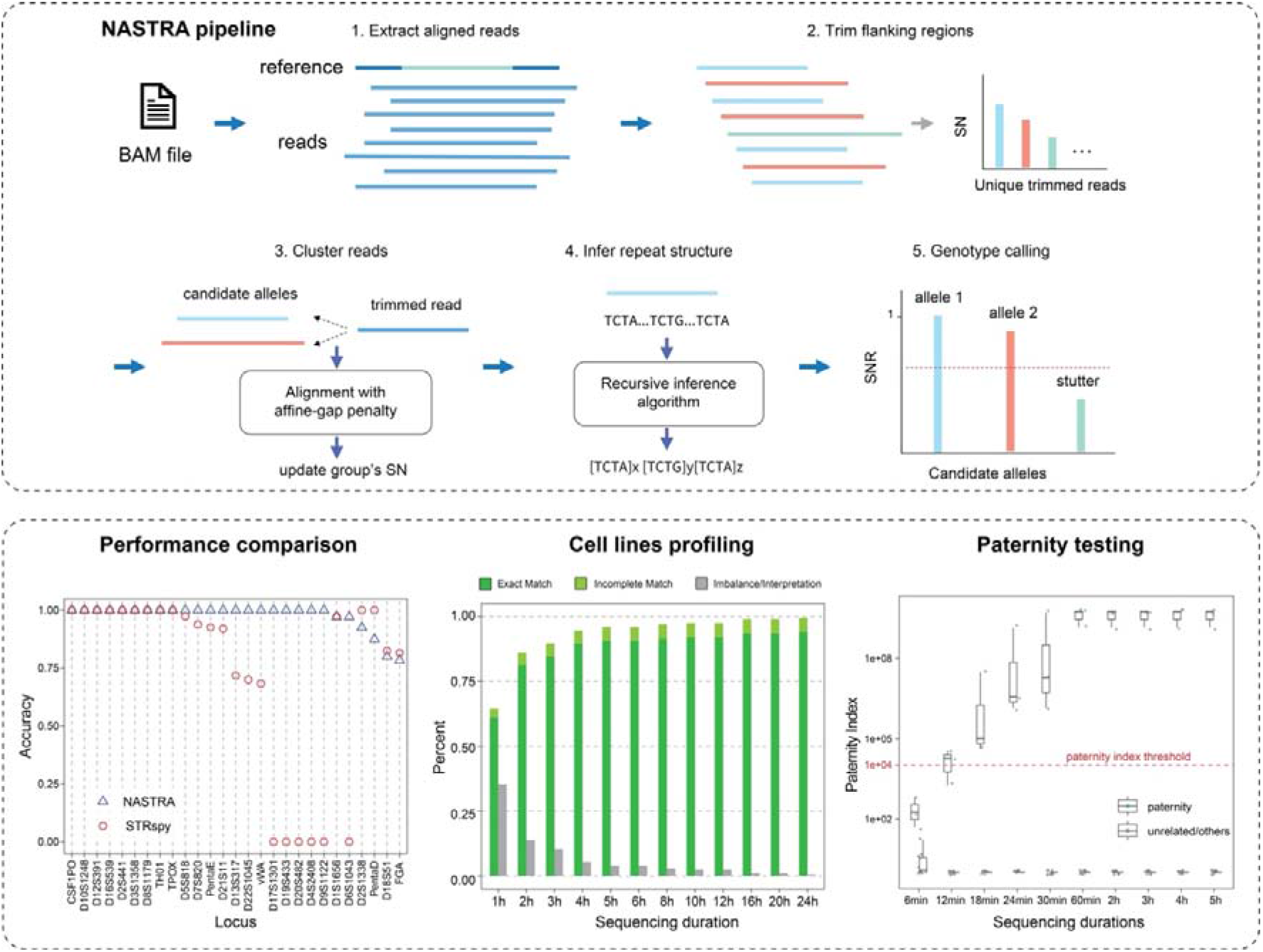

## INTRODUCTION

Short tandem repeats (STRs) in the human genome are specific DNA loci that can have a high degree of polymorphisms in the number and forms of repeat motifs among individuals. This feature makes STRs a kind of genetic markers well-suited for individual identification (1, 2), cell line authentication (3, 4), and relationship inferring (5–7). STR genetic markers are usually selected considering high heterozygosity, regular repeat units, distinguishable alleles, and robust PCR amplification. The Short Tandem Repeat DNA Internet Database (STRBase) maintained by the National Institute of Standards and Technology (NIST) (8) includes more than 70 human STR genetic markers, with numbers of known per-locus alleles ranging from 19 (D10S1248) to 114 (FGA). These STRs have been utilized for decades by forensic laboratories world-wide and have formed large-scale human DNA databases for public security purposes in many countries. In biomedical research, STR profiling is used for cell-line authentication, which plays an important role in ensuring scientific reproducibility and preventing misidentification as well as cross-contamination of cell lines (9, 10). The International Cell Line Authentication Committee (ICLAC, https://iclac.org) integrates many online databases and search tools, such as the Cellosaurus knowledge resource (11), DSMZ STR profile database (12), CLIMA database (13), and CLASTR (14).

The traditional approach to STR profiling relies on capillary electrophoresis (CE)-based length analysis. Massively parallel sequencing (MPS) has also been validated for use in STR profiling, offering the capability to analyze a broader array of STRs in a single run compared to the CE method. However, both CE and MPS systems for STR profiling have their limitations: they are immobile, expensive, and require specific environmental conditions, such as stable temperature and absence of vibration. Consequently, these instruments are typically housed in well-equipped laboratories or specialized facilities. This necessitates the transportation of samples for analysis, often leading to delays of several days before obtaining STR typing reports.

Nanopore sequencing offers an economical and flexible alternative for STR profiling. Devices like the MinION Mk1B/Mk1c (Oxford Nanopore Technologies, Oxford, UK) and QNome-3841 (Qitan Technology, Chengdu, China) stand out for their exceptional portability and lower cost compared to CE and MPS systems. These portable nanopore sequencing devices not only produce longer reads than Sanger sequencing and NGS systems but also yield data at the Gbp level in a single run. Nanopore sequencing has found diverse applications, including on-site genomic surveillance of pathogens (15, 16) and biodiversity studies (17, 18). Zaaijer et al. (19) conducted a pilot study using the MinION device for rapid whole genome sequencing to authenticate human cell lines in biological research.

Despite these advantages, nanopore sequencing is noisier than CE and MPS, which is challenging for the accurate analysis of the STR markers (20–23). For the detection of kilo-nucleotide long STRs in repeat expansion disorders, tools like repeatHMM (24), STRique (25), and DeepRepeat (26) have been developed. Compared to detecting abnormally long expansions, conducting accurate genotyping calls for cell line identification and forensic applications presents greater challenges. The alleles of these STR genetic markers typically range from 100 to 300 nucleotides (nt) in length and are composed of 3- to 5-nt repeat units. The genotypes of these STR markers are determined by the precise count of their repeat units. We previously reported that tools designed for detecting repeat expansions, like repeatHMM (24), are not suitable for STR genetic markers. In contrast, employing the classic Smith-Waterman algorithm for re-alignment significantly enhances the accuracy of STR typing, as noted in our earlier findings (27). Hall et al. (23) and Tytgat et al. (21) and recently Lang et al (28), have employed a straightforward strategy. Their approach involves aligning all available sequences of STR alleles and selecting the alleles with the most supporting reads as the genotypes. However, this method relies on allele databases, which can lead to typing errors for carriers of unknown alleles. In the meantime, owing to the inherent noise in nanopore sequencing, there’s a heightened risk of errors when using alignment-based strategies for alleles with highly similar nucleotide sequences.

To address the precision challenges in STR typing using nanopore sequencing, we introduced NASTRA, an algorithm specifically developed for accurate STR analysis without reliance on reference alignment. It’s particularly suited for cell line authentication, as well as individual and kinship identification. NASTRA utilizes read clustering to reduce the impact of subtle sequencing errors and features a recursive algorithm for inferring the repeat structure of alleles based solely on known repeat motifs. Our evaluation, encompassing various multiplex systems and flow cells, demonstrated NASTRA’s effectiveness in genotyping 76 individual samples across 27 autosomal STR markers. Additionally, genotyping calls on 8 standard cell lines resulted in 100% accuracy. We further validated NASTRA in individual identification and paternity testing within a four-member family and two unrelated individuals. The results demonstrated that NASTRA could accurately confirm all individual identifications and paternity tests within sequencing durations of 12 and 18 minutes, respectively. Overall, NASTRA exhibits impressive performance in both accuracy and speed.

## MATERIAL AND METHODS

### Sample collection and DNA extraction

A total of 76 blood samples were obtained from unrelated, anonymized Han Chinese volunteers, comprising 38 females (ID: F01-F38) and 38 males (ID: M01-M38). Of these, 30 samples (F01-F15 and M01-M15) were sourced from our previous study(27), while the remaining 46 samples (F16-F38 and M16-M38) were newly acquired. Besides these, we also gathered 8 control DNA samples. These included 3 forensic DNA controls (2800M from Promega, 9947A, and 9948 from Origene), along with 5 cell line standards (NA12878, NA24143, NA24149, NA24694, and NA24695 from the Coriell Institute). Additionally, for a case study on paternity testing, we included a family of four: a grandmother, mother, father, and child. Genomic DNA from the newly collected samples was extracted using the PureLink Genomic DNA Kit (Invitrogen, USA). Prior to amplification, all DNA samples were quantified using the Qubit 3.0 Fluorometer (Invitrogen) and the Qubit dsDNA HS Assay Kit, following the manufacturer’s instructions.

### Benchmarking data using MiSeq FGx system

We amplified the newly collected DNA samples using the ForenSeq DNA Signature Prep Kit (DNA Primer Mix A). Concurrently, the 2800M sample was amplified as a positive control using DNA Primer Mix B, encompassing all STR loci present in the Mix A kit. To ensure ample amplicons for both Illumina MiSeq FGx and Nanopore sequencing, we used 50 ng of template DNA instead of the recommended 1 ng. The PCR amplifications were executed on the ProFlex™ 3×32-Well PCR System (Thermo Fisher). Subsequently, all purified amplified libraries were evenly split into two portions for sequencing on the MiSeq FGx system and Nanopore platform. The pooled libraries were prepared by combining equal volumes (2 μL) of each normalized library (10 nmol), which were then diluted to a final concentration of 4 nmol. For MiSeq FGx sequencing, 7 pmol of the pooled libraries were used. All experimental procedures were meticulously conducted in accordance with the manufacturer’s guidelines. Genotype analysis of the STR loci was conducted using the ForenSeq Universal Analysis Software, UAS (v1.3.6897; Verogen) with default parameters.

### Nanopore sequencing on ForenSeq amplicons

Libraries were prepared with the Ligation Sequencing Kit (SQK-LSK109). The process involved processing 0.2 pmol of purified DNA per sample using the NEBNext Ultra II End repair/dA-tailing Module (E7546). Samples were then multiplexed utilizing both the Native Barcoding Expansion 1-12 (PCR-free) and 13-24 (PCR-free) kits. Subsequently, adapters were ligated to the pooled libraries using the NEBNext Quick Ligation Module (E6056). A total of 0.05 pmol of these libraries were loaded on the R9.4.1 flow cells. The sequencing was carried out using MinKNOW software. Forty-six DNA samples, along with eight control DNA samples, underwent sequencing across three runs, each lasting over 24 hours. For the paternity testing, six samples were loaded on an R10.3 flow cell for sequencing.

### Nanopore sequencing for PowerSeq amplicons

Following the manufacturer’s guidelines, we selected 48 DNA samples for amplification using the PowerSeq™ 46GY System (Promega, WI, USA). We utilized 1 ng of template DNA per sample and conducted PCR amplification over 29 cycles. Then, the PCR products were purified using the AmPure XP Beads (Beckman Coulter, Indianapolis, USA), and the DNA concentrations were quantified. The library preparation process mirrored the methodology previously described. The final step involved loading 0.05 pmol of the libraries on the R10.3 flow cells for sequencing, which was executed using MinKNOW software. In this phase, the 48 DNA samples and the 6 samples involved in paternity testing case study underwent sequencing across 3 runs, each extending beyond a 24-hour duration.

### Capillary Electrophoresis for paternity testing

For the paternity testing case study, we performed STR typing on a family of four individuals and two unrelated individuals, 2800M and 9948. The typing results were obtained by capillary electrophoresis and served as a benchmark. We used 1 μl of DNA template from each sample to amplify the targeted amplicons with the PowerPlex 21 kit (Promega) and the GeneAmp PCR System 9700 (Applied Biosystems). The PCR products were then separated using the 3130XL Series Genetic Analyzers (Applied Biosystems). Data analysis was conducted using the GeneMapper ID Software (version 3.2) (Applied Biosystems).

### Bioinformatic analysis for nanopore sequencing data

The basecalling process was performed using Guppy v6.3.4 with the high-accuracy model. Then, the reads were aligned to the human reference genome (assembly GRCh37, hg19) using Minimap2 v2.17-r941 (29, 30). Finally, SAMtools v1.6 (31) were employed to convert SAM files into BAM format, which were then used as input for NASTRA. Down sampled data were generated using NanoTime, which is available at https://github.com/renzilin/NanoTime. In this process, NanoTime utilized the timestamps of each read in the sequencing summary file to generate a series of FASTQ files with varying sequencing durations ranging from 1 to 24 hours (1, 2, 3, 4, 5, 6, 8, 10, 12, 16, 20, and 24 hours).

## RESULTS

### Overview of NASTRA

NASTRA requires three inputs: (1) the alignments of nanopore reads to the human reference genome; (2) the reference genomic positions of STR markers; and (3) the repeat motifs stored in the STR fact sheets. NASTRA’s pipeline consists of two primary components: clustering of reads and repeat structure inference. These two components are designed to mitigate the potential impact of subtle sequencing errors and make accurate genotyping calls without the allele reference database (Figure 1b).

**Figure 1.**
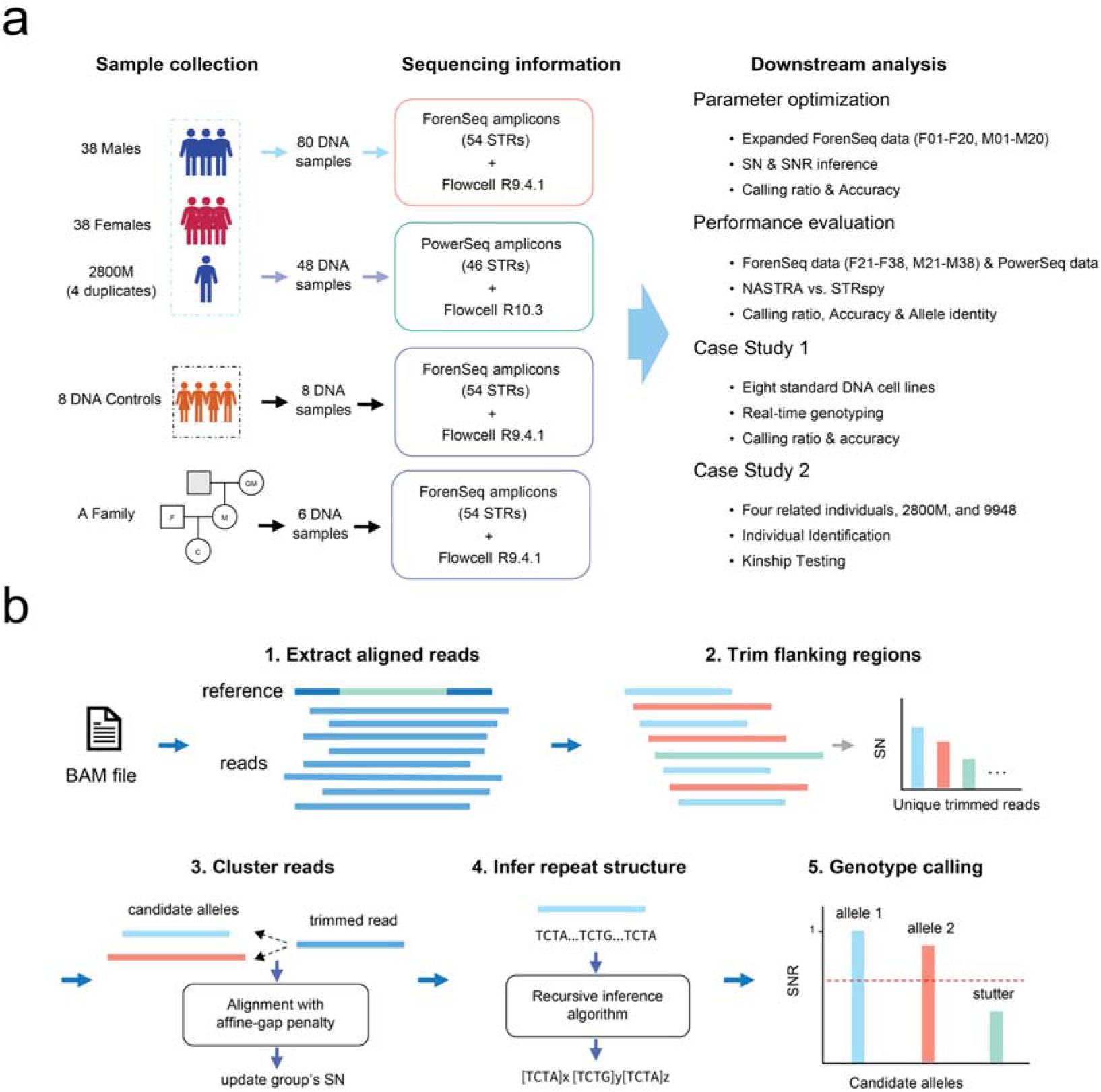
The overview of the study design. (**a**) We carried out nanopore sequencing on a collection of 76 individual samples, along with duplicated 2800Ms, using various amplification kits and flow cells. This was done for both parameter optimization and performance evaluation. Subsequently, we performed two case studies, one involving cell line profiling and the other focused on paternity testing. (**b**) The framework of NASTRA. NASTRA consists of five steps to perform genotyping calls.

In step 1, NASTRA retrieves aligned reads for a target locus from the BAM file using the positional information of the locus. In step 2, NASTRA employs local pairwise alignment with an affine-gap penalty to trim the prefix and suffix flanking sequences, with an extra three bases remaining on each side. As a result, each unique type of these trimmed reads is grouped, with their read counts being aggregated. Due to the presence of subtle sequencing errors, the number of these groups may exceed 100. To address this, NASTRA uses global pairwise alignment with an affine-gap penalty to cluster the trimmed reads in step 3, tolerating minor sequencing errors like single-base InDels or SNPs. Subsequently, the read counts of these clustered groups are summed up, referred to as the Supporting Read Number (SN).

The top three sequences with the highest SN are selected as candidate alleles. In step 4, we have developed a recursive algorithm that infers the repeat structure of each allele sequence based on repeat motifs. This approach not only facilitates the rapid acquisition of STR genotypes but also assists in the quick identification of Single Nucleotide Variants (SNVs) in genotypes (Figure 2). Ultimately, NASTRA carries out STR genotyping by utilizing the SN and the Supporting Read Number Ratio (SNR), where SNR is calculated as the ratio of an allele’s SN to the SN of the major allele. Drawing inspiration from the quality control mechanism used in the ForenSeq Universal Analysis Software (UAS), NASTRA incorporates similar rules. These rules are designed to filter out unreliable genotyping results characterized by low coverage and stutters, thereby enabling more accurate genotyping calls.

**Figure 2.**
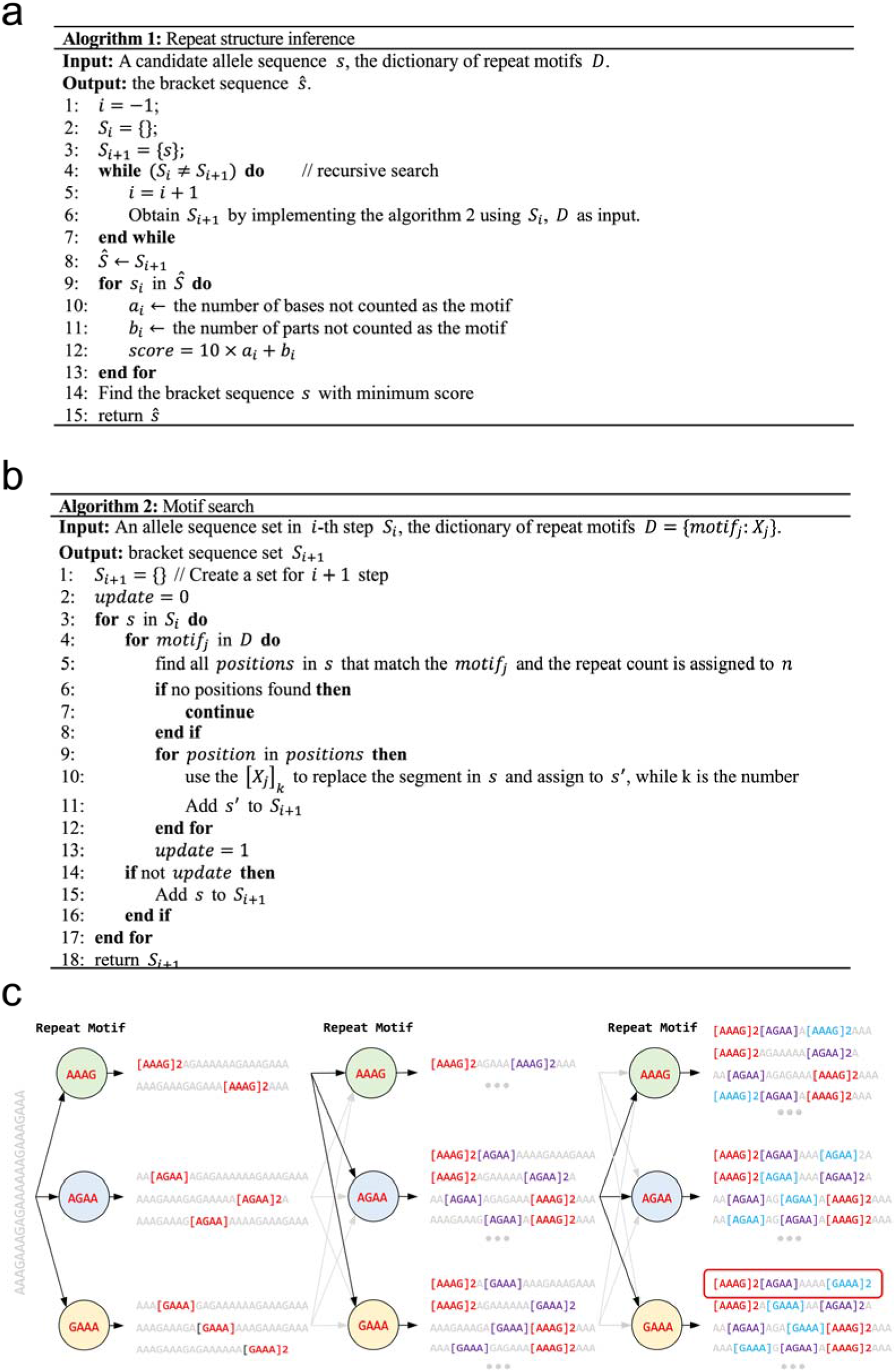
The recursive algorithm for repeat structure inference. (**a**) Pseudocode for the repeat structure inference. This algorithm utilizes the motif search method described in (**b**), employing recursion to achieve the structure inference. (**b**) Pseudocode for the motif search algorithm. (**c**) This schematic illustrates the process of inferring repeat structure. The sequence depicted in gray on the left represents the candidate allele. In each step, motifs, AAAG, AGAA, and GAAA, are searched one by one. As the process progresses, identified fragments in each step are transformed into bracket sequences, which are highlighted in red.

For the pairwise alignment used in NASTRA, we used parasail-python (32) (https://github.com/jeffdaily/parasail-python) to conduct pairwise alignment with an affine-gap penalty in trimming and clustering steps. The parameters were set with a match score of 75, a mismatch penalty of 90, a gap opening penalty of 75, and a gap extension penalty of 10. As a result, this alignment configuration shows a tendency to prefer two consecutive gaps over a single mismatch. However, if the sequence has more than two continuous gaps, the alignment outcome becomes more inclined to favor a single mismatch.

### Repeat structure inference and genotyping calls

Repeat motifs for each STR were collected from the STR fact sheet of STRBase (8). The algorithm is designed to recursively search for sequence fragments that match these repeat motifs. To elucidate the process, we provided pseudocode (Figure 2a-b) and an example (Figure 2c). NASTRA begins by searching for each motif (AAAG, AGAA, and GAAA) within a given sequence (depicted in full grey). After finding these motifs, it identifies 8 unique fragments and transforms them into 8 distinctively marked sequences. The algorithm then repeats the search for each marked sequence. This iterative process is carried out until no new fragments are found, or the iteration count exceeds 50. Finally, NASTRA calculates the count of non-motif bases and parts for each marked sequence. The sequence with the lowest score (defined in Figure 2a) is then determined to be the most plausible inferred structure (highlighted in a red box). For each structured allele, NASTRA sums up the counts of repeat motifs to infer its genotype. However, the presence of certain single nucleotide polymorphisms (SNPs) in the adjacent and repeat regions (33) can lead to challenges in making accurate genotyping calls. To address this, locus-specific adjustments have been incorporated into NASTRA. Detailed information about these adjustments is available in the Supplementary Materials.

NASTRA determines the STR’s genotype based on the alleles’ SNs and SNRs. In more detail, if the highest SN among the alleles is less than 10, NASTRA will categorize the result as ‘Interpretation’, indicating an interpretation failure due to inadequate coverage. Meantime, NASTRA assesses whether the STR genotype is heterozygous or homozygous based on the SNR of the allele with the second-largest SN. A heterozygous result is declared if the SNR of the minor allele surpasses a predefined threshold. Conversely, if the SN of the minor allele falls below this threshold, but the major allele’s SN exceeds it, the result is labeled as ‘imbalance’. For the optimal selection of SN and SNR thresholds, we have included a configuration file for users.

### Parameter optimization using down sampled data

To infer the optimal parameters for NASTRA, we split the samples into two sets: (1) training set, comprising 20 males and 20 females, and (2) test set, consisting of 18 males, 18 females, and 4 replicates of 2800M samples. Considering the limited size of the training set and the insensitivity of the deep sequencing depths toward the thresholds, we expand the training and test sets using the tool NanoTime. To assess the performance of NASTRA, we used the benchmark set built by the MiSeq FGx system, where the typing results with “imbalanced,” “allele count,” “stutter,” and/or “interpretation threshold” were excluded.

First, we used NASTRA to genotype alleles for 12, 960 STRs on 27 autosomal STR loci across 480 samples. Then, we used various SN values (ranging from 0 to 50 in increments of 5) as well as SNR values (ranging from 0 to 1 in increments of 0.01) to accomplish the genotyping calls for these STRs. The genotyping results with “Interpretation” and “Imbalance” were removed. Finally, we compared the NASTRA’s genotyping results to the benchmark set, and the results were classified into five categories: (1) **Exact Match**: correct genotyping; (2) **Incomplete Match**: correct genotyping when the incomplete repeat unit is ignored; (3) **One Match**: one of the alleles is genotyped correctly; (4) **Incomplete One Match**: one of the alleles is correctly genotyped, when the incomplete repeat unit is ignored; (5) **Mismatch**: incorrect genotyping.

For each SN value, we calculated the maximum genotyping accuracy for each autosomal STR locus on the expanded training set. The result showed that when SN reaches or exceeds 25, the maximum genotyping accuracy for all STR loci can exceed 95% (Figure 3a). After having determined the SN value of 25, we calculated the genotyping accuracy for each autosomal STR across different SNR values. The SNR threshold is defined as the median value within the range of SNR values corresponding to the peak genotyping accuracy (Figure 3b). After having determined the threshold SN and SNR, NASTRA was used to perform STR genotype calling on the expanded test set.

**Figure 3.**
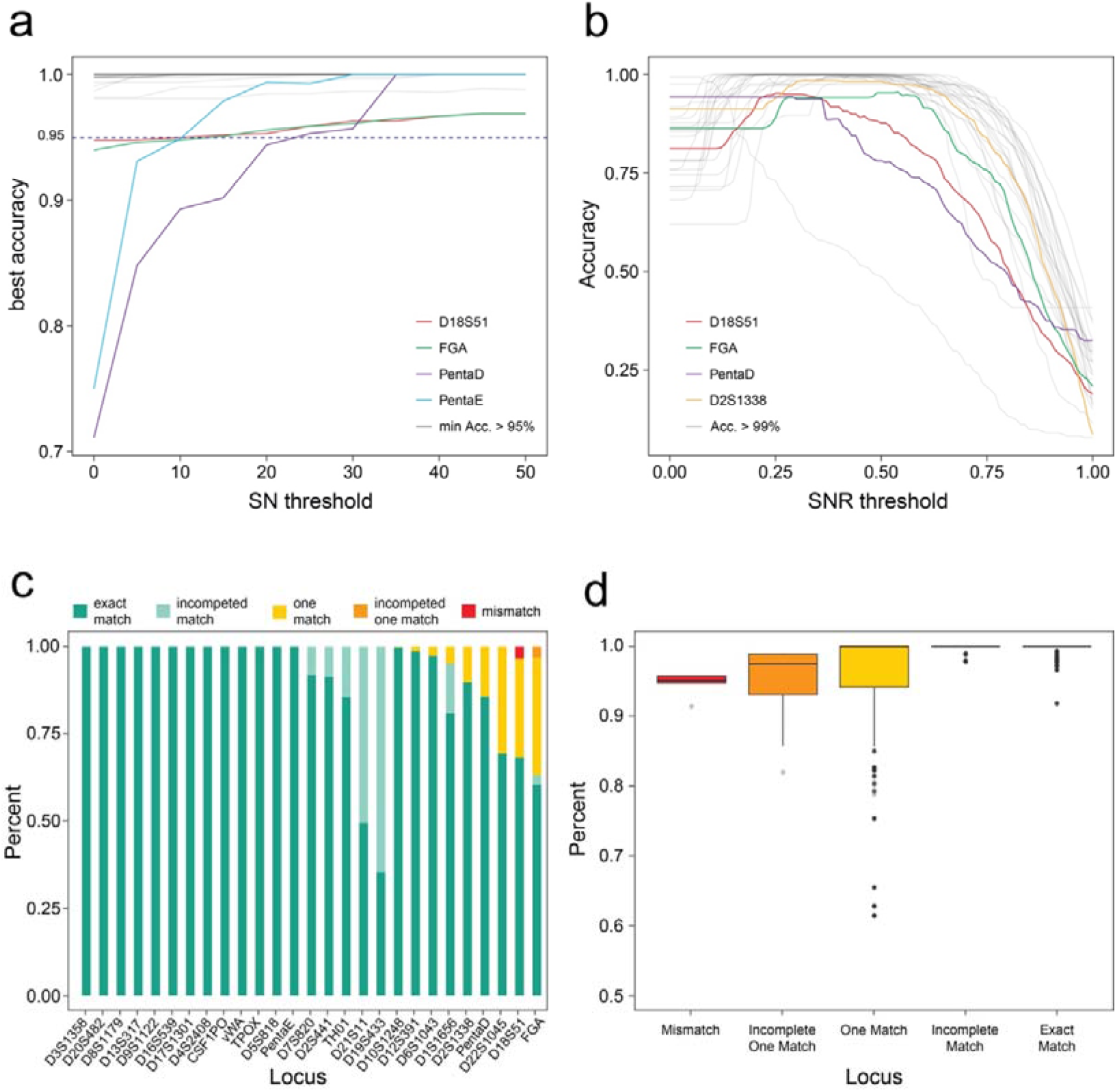
Parameter optimization using downsampled data. (**a**) NASTRA’s highest accuracy on each locus with a fixed supporting read number (SN). Each line represents a locus, and the gray lines are loci whose typing accuracy is always greater than 95%. (**b**) The genotype calling accuracy of NASTRA on each locus at various supporting read number ratios (SNR), with the SN fixed at 25. Each line represents a locus, and the gray lines are loci whose peak typing accuracy surpasses 99%. (**c**) The statistical diagram presents the genotyping results from downsampled data. (**d**) The blast identity of allele sequences predicted by NASTRA with the benchmark sequences. In (**c**) and (**d**) dark green indicates an exact match, light green signifies an incomplete match, yellow for one match, orange for an incomplete one match, and red denotes a mismatch.

In summary, 3, 532 of 12, 960 genotype calls failed to pass the quality control, including 984 calls in the benchmark set and 2, 548 calls by NASTRA, which is mainly caused by insufficient calls with short sequencing duration (Supplementary Figure S1). The rest 9, 428 of 12, 960 (72.8%) STRs on 480 expanded test sets were genotyped successfully, consisting of 8, 455 (89.7%) exact matches, 642 (6.8%) incomplete matches, 307 (3.3%) one matches, 11 (0.1%) incomplete matches, and 13 (0.1%) mismatches. Figure 3c depicts the accuracy of each STR across the 480 samples. Out of the 27 loci, 18 loci had a genotyping accuracy of 100% (exact match or incomplete match), and 4 loci had an accuracy greater than 95%. Additionally, we employed pairwise alignment to investigate sequence identity for each allele (Figure 3d). We found that for most “incomplete match” genotyping results, the sequence identity of the allele sequences is equal to 1. This suggested that NASTRA preserved the true allele sequence, while solely disregarding incomplete repeat units in terms of repeat counting.

### Evaluation of NASTRA on ForenSeq amplicons with R9.4.1 flow cells

We assessed the performance of NASTRA on ForenSeq amplicon sequencing data with a sequencing duration of 24 hours, comparing it to the alignment-based method, STRspy(23). STRspy is an STR profiling tool tailored for long-read sequencing, which aligns reads to the allele reference database. STRspy provides a customized database that contains 638 allele sequences of 22 autosomal STRs. Thus, there are 5 STR loci (D17S1301, D20S482, D4S2408, D9S1122, and D6S1043) that can’t be genotyped due to the absence of allele information. We first used the calling ratio, the proportion of samples successfully genotyped at a locus, to assess both tools. There are 22 of 27 loci exhibiting a calling ratio of 1 using STRspy. Regarding NASTRA, the calling ratios for 22 out of the 27 loci range from 92.1% to 100%, as illustrated in Supplementary Figure 2. For genotyping accuracy, NASTRA demonstrated superior performance compared to STRspy. NASTRA genotyped accurately 21 loci across all samples, while STRspy successfully genotyped 11 loci correctly (Figure 4a, Supplementary Table S1). For NASTRA, the deviation between true alleles and inferred alleles is mainly attributed to the incomplete repeat (deviation < 1, Figure 4b). While using STRspy, the absence of some specific allele information influenced the genotyping accuracy. For instance, the FGA’s allele 24.2 of sample F25 is [GGAA]2_GGAG_AA_[AAAG]16_[AGAA]1_AAAA_ [GAAA]3, while the allele in the reference database is in the [GGAA]2_GGAG_[AAAG]17_AA_AAAA_[GAAA]3. Thus, STRspy predicted the allele as 24, leading to an incomplete match with the wrong repeat structure. This highlights the importance of a comprehensive and highly reliable reference database for alignment-based methods to achieve precise genotyping.

**Figure 4.**
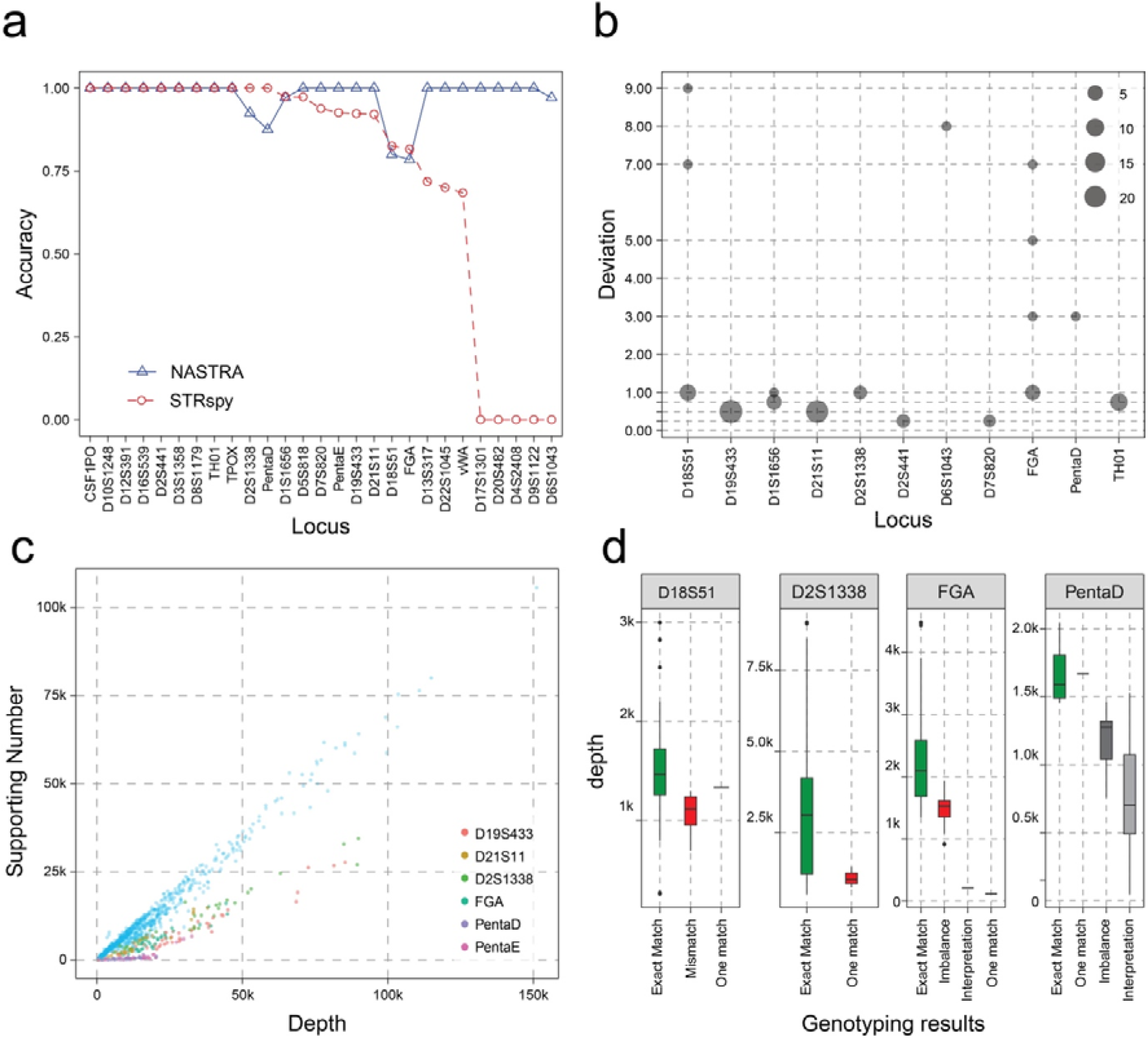
Performance evaluation on ForenSeq data with R9.4.1 flow cells. (**a**) Accuracy comparison between NASTRA (blue) and STRspy (red). (**b**) Genotype deviations from baseline as calculated by NASTRA. For heterozygous genotypes, the sum of the deviation for both alleles was calculated. (**c**) The correlation of SN and sequencing depth. Blue dots represent other 21 autosomal loci. (**d**) Distribution of sequencing depth across various genotyping results at four loci. Green indicates exact match, red denotes mismatch, dark gray represents imbalance, and light gray signifies interpretation.

To investigate the causes of incorrect genotyping results, we analyzed the relationship between SN and sequencing depth. Figure 4c indicates that the loci with incorrect genotyping have lower SNs than those with correct genotyping when the sequence depth is close. This phenomenon may be attributed to the presence of homopolymers containing adenine (A) and guanine (G), as observed in loci such as D18S51, FGA, PentaD, and D2S1338. A possible solution is to increase the sequencing depth for these loci (Figure 4d).

### Evaluation of NASTRA on PowerSeq amplicons with R10.3 flow cells

To assess the performance of NASTRA on different amplification systems and flow cells, we conducted a sequencing experiment on 46 samples using Promega’s PowerSeq 46GY kit and R10.3 sequencing flow cells. The PowerSeq kit is a co-amplification system consisting of 22 autosomal STR loci and 23 Y-STR loci. The ONT R10.3 flow cell represents a new version featuring a new pore design, resulting in enhanced read accuracy and quality.

Using the optimal parameters inferred on ForenSeq amplicons with R9.4.1 flow cells, NASTRA showed the robustness of accuracy on the PowerSeq data (Figure 5a). For NASTRA, among the 22 autosomal STR loci, 20 STR loci exhibited a calling ratio of 1 (Supplementary Figure 3), with only the remaining 2 loci, PentaE (94.7%) and PentaD (55.6%). For genotyping accuracy, D18S51, FGA, and PentaD were still not typed correctly (Figure 5b). Additionally, new loci with incorrect genotyping results (TH01 and PentaE) were found in the PowerSeq data, possibly attributed to the overall shorter amplicons in the PowerSeq data compared to the ForenSeq data (Supplementary Figure 4). Despite this issue, NASTRA demonstrated superior performance over STRspy, with 17 out of 21 loci correctly typed compared to 11 (Supplementary Table S2). Specifically, NASTRA achieved an accuracy of over 95% in 18 loci, compared to STRspy’s 13 loci. The accuracy of the remaining 4 loci ranged from 80% to 91.5%. Meanwhile, STRspy achieved an accuracy between 61.7% and 91.3% in the remaining 7 loci. In Figure 5c, the primary deviation in most genotyping calls, apart from the loci D18S51, PentaD, and PentaE, is attributed to the presence of incomplete units.

**Figure 5.**
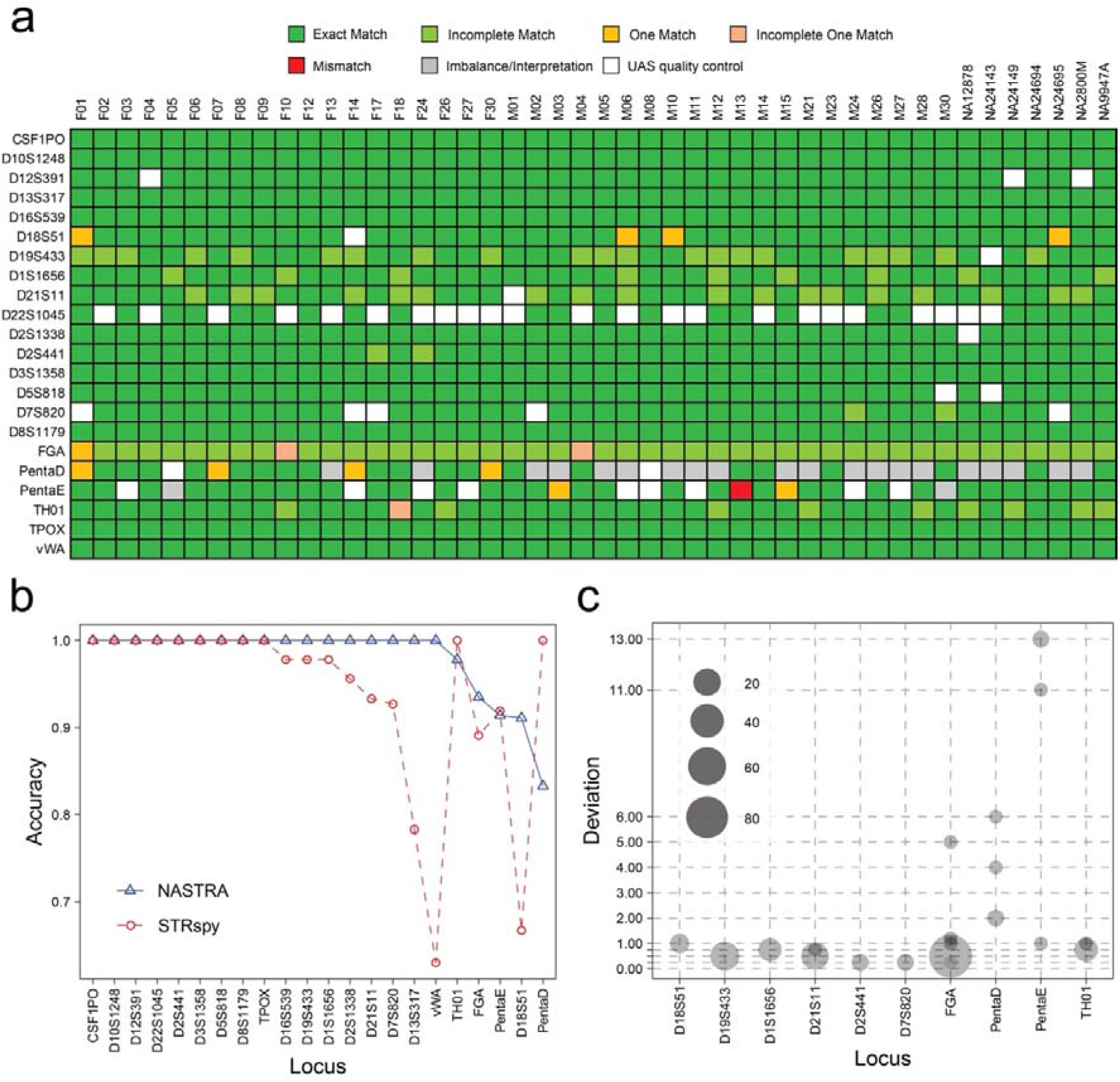
Performance evaluation on PowerSeq data with R10.3 flow cells. (**a**) Overview of genotyping results on 22 autosomal STRs across 46 samples. (**b**) Accuracy comparison between NASTRA (blue) and STRspy (red). (**c**) Genotype deviations from baseline as calculated by NASTRA. For heterozygous genotypes, the sum of the deviation for both alleles was calculated.

### Case study: real-time typing of 8 human cell lines

We evaluated NASTRA for cell line authentication by profiling the STRs in eight individual cell lines. For this, we processed DNA samples using the ForenSeq Signature kit and then performed sequencing on a MinION platform with an R9.4.1 flow cell. The genotyping results with different sequencing durations are shown in Figure 6a. Excluding unreliable genotyping calls (shown in grey) and those that failed UAS quality control in the benchmark (shown in white), NASTRA accurately genotyped all samples. When the sequencing time extended to four hours, the CODIS core loci in all individuals were genotyped correctly, demonstrating NASTRA’s sufficiency for individual identification and cell line authentication. Notably, our analysis solely concentrated on autosomal STRs, which means a portion of the sequencing throughput of other markers (24 Y-STRs, 7 X-STRs, and 94 identity SNPs) was underutilized. This suggests that the sequencing duration required for cell line authentication could be shorter. We found that the unreliable genotyping results were mainly caused by the sequencing depth, as shown in Figure 6b. When the sequencing depth for each locus exceeds 500, the number of unreliable genotyping results would be very few (Figure 6c). This would provide a certain reference for the design of the multiplex panels tailored for nanopore sequencing.

**Figure 6.**
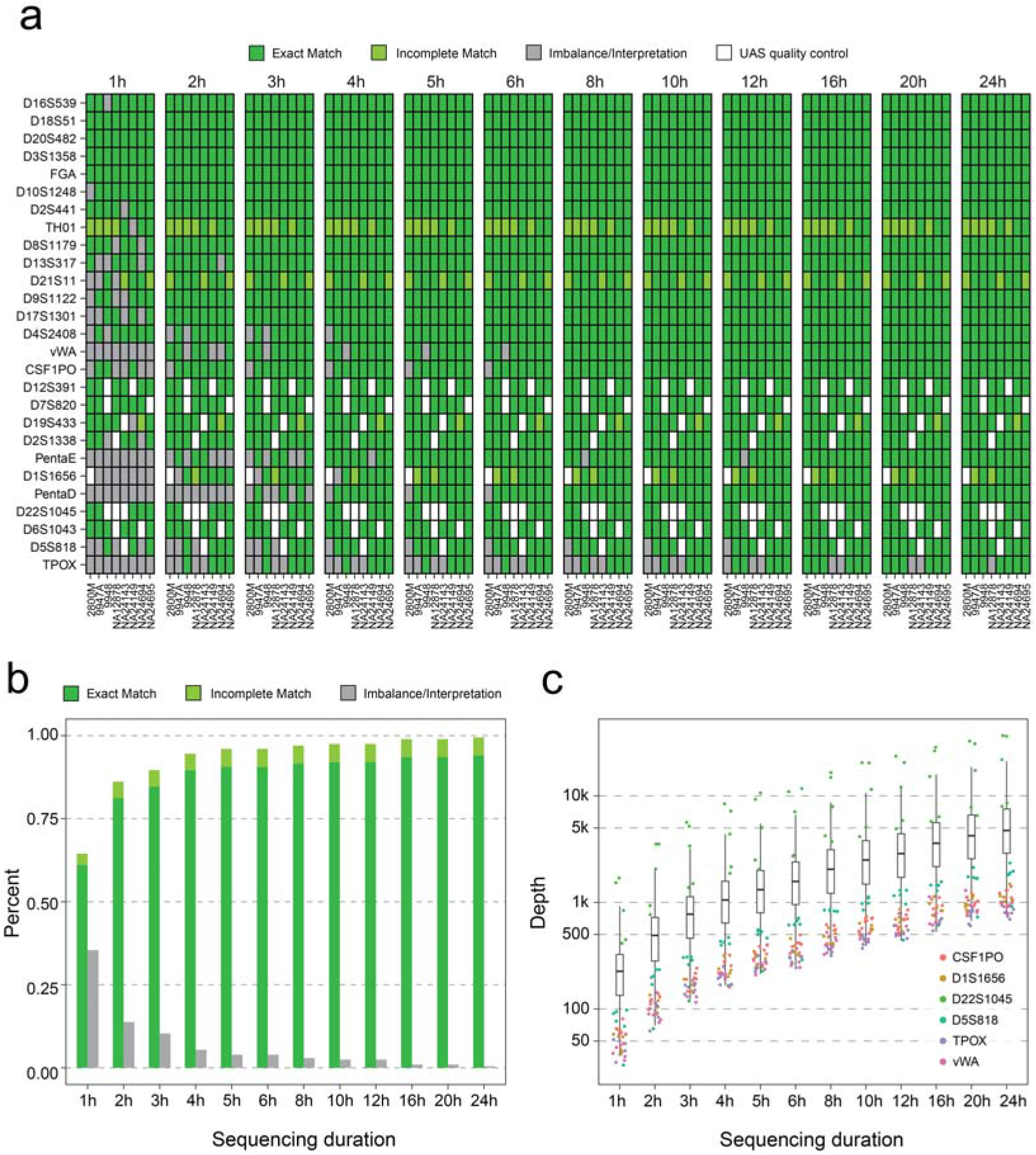
Performance evaluation of profiling 8 standard cell lines. (**a**) Overview of genotyping results of 8 standard DNA samples across 27 autosomal STRs with different sequencing durations using an R9.4.1 flow cell. Grey represents unreliable results and white represents genotypes that did not pass UAS quality control in the benchmark construction. (**b**) The statistical diagram presents the proportion of genotyping results at various sequencing durations. (**c**) Distribution of sequencing depth at 27 autosomal STR loci over different sequencing times. D22S1045 shows the highest sequencing depth, whereas CSF1PO and four other loci consistently show low sequencing depths.

### Case study: paternity testing on a family of four individuals

To further evaluate NASTRA, we carried out a paternity test on a family of four. Additionally, we included two unrelated DNA samples, 2800M and 9948, as controls (see Figure 7a). The benchmark for the test was established using capillary electrophoresis. For nanopore sequencing, we utilized the ForenSeq Signature kit to generate amplicons and employed an R10.3 flow cell for the sequencing. We first summarized the sequencing throughput at different durations (Supplementary Table S3). On the individual level, the MinION yielded sequencing data ranging from 1.84 Mbp (M, 6 minutes) to 95.50 Mbp (F, 5 hours). We then assessed NASTRA’s genotyping accuracy on 21 STR loci among 6 individuals (122 genotyping results, Figure 7b). When the sequencing duration was 6 minutes, the majority of genotyping results did not meet quality control standards, primarily due to inadequate sequencing yields. When the sequencing duration reached 60 minutes, all STR loci were correctly genotyped. To explore the sequencing duration required for individual identification and paternity testing, we calculated the likelihood ratio for individual identification and the paternity index for individual pairs. With a 12-minute sequencing, a minimum of 11 loci were genotyped across the six individuals (2800M, 4.27 Mbp), achieving a likelihood ratio surpassing 1e15 (Figure 7c). According to the standards, a parent-child relationship is confirmable with a paternity index over 1e4. At 18 minutes, three parent-child relationships were successfully confirmed (Figure 7d). It’s important to note that a considerable portion of the sequencing data, including SNPs and sexual STRs amplicons, was not used in our analysis due to the ForenSeq system. This suggests that the duration for both identification tasks could be potentially reduced.

**Figure 7.**
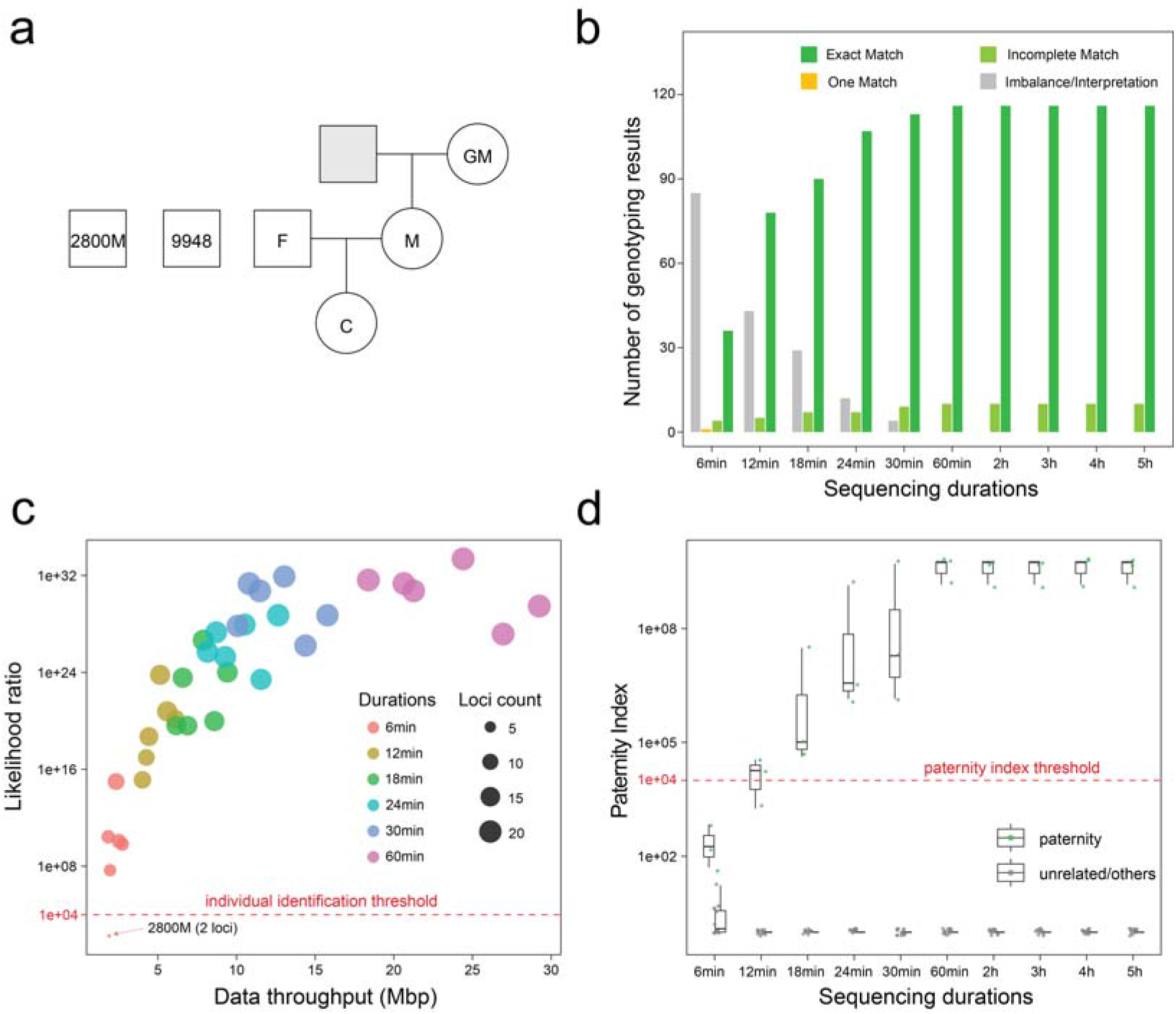
A case study of using NASTRA to perform individual identification and paternity testing. (**a**) Sample and family information. (**b**) Counts for different genotyping results with varying sequencing durations. (**c**) Likelihood ratios for individual identification. X-axis represents the sequencing throughput of ForenSeq data for each individual. The size of the dots represents the number of loci successfully and correctly genotyped. Colors represent for sequencing duration, and red dashed line represents for the individual identification threshold. (**d**) Distribution of paternity index scores for parent-child paternity testing with various sequencing durations. Green points represent parent-child relationships, while gray points represent unrelated pairs.

### Comparison of running time

NASTRA is developed using Python programming language and integrates the parasail library (32), which is a C-based pairwise alignment tool. When using NASTRA, users can accomplish parallel computing at the individual level by invoking the ‘xargs’ shell command. For STRspy, the developers utilize the ‘parallel’ shell command for parallelization at the locus-level. To conduct the runtime comparisons, we allocated the same number of threads (eight) to both NASTRA and STRspy. The STR genotype calling for all sequencing data was conducted on a server equipped with a 64-Core/128-Thread AMD EPYC 7763 Processor (2.45GHz) and 256 GB of memory.

Table 1 presents a detailed runtime usage between NASTRA and STRspy. It reveals that NASTRA is approximately 60 times faster than STRspy when analyzing ForenSeq amplicon sequencing data. The PowerSeq data had more reads to process than the ForenSeq data because a smaller fraction of the sequencing throughput was invalidated. This resulted in a longer alignment runtime. Consequently, NASTRA demonstrates a significantly faster performance overall. However, the two tools are developed in different programming languages, making direct runtime comparisons somewhat subjective. Notably, our analysis shows that STRspy’s time-consuming alignment process, which includes individual executions of Minimap2 (29, 30) for each locus, contributes substantially to its longer runtime.

**Table 1.**
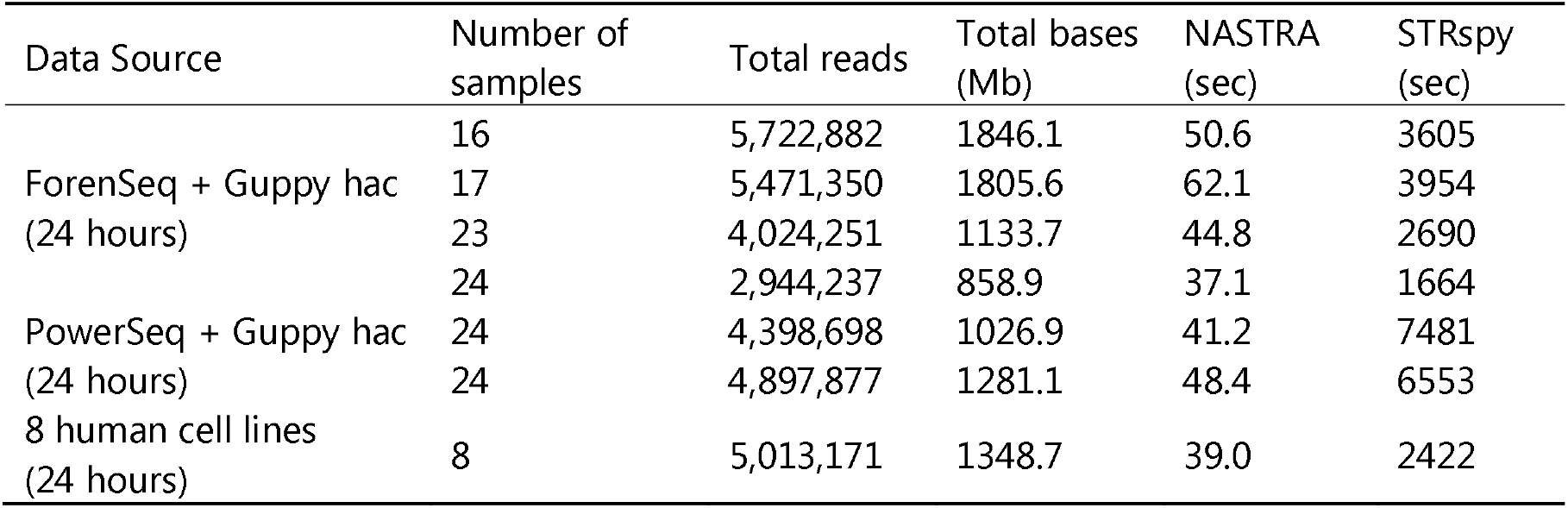
Runtime comparison under the same thread count (8 threads).

| Data Source | Number of samples | Total reads | Total bases (Mb) | NASTRA (sec) | STRspy (sec) |
| --- | --- | --- | --- | --- | --- |
| ForenSeq + Guppy hac (24 hours) | 16 | 5,722,882 | 1846.1 | 50.6 | 3605 |
|  | 17 | 5,471,350 | 1805.6 | 62.1 | 3954 |
|  | 23 | 4,024,251 | 1133.7 | 44.8 | 2690 |
|  | 24 | 2,944,237 | 858.9 | 37.1 | 1664 |
| PowerSeq + Guppy hac (24 hours) | 24 | 4,398,698 | 1026.9 | 41.2 | 7481 |
|  | 24 | 4,897,877 | 1281.1 | 48.4 | 6553 |
| 8 human cell lines (24 hours) | 8 | 5,013,171 | 1348.7 | 39.0 | 2422 |

## DISCUSSION

In this study, we introduced NASTRA, a novel computational approach designed for STR analysis. NASTRA is built on two primary modules: read clustering and repeat structure inference. To mitigate the influences of sequencing noises on genotyping calls, NASTRA employs read clustering. This approach identifies candidate alleles by discounting minor noises. Following this, the structure-aware algorithm reconstructs the repeat structures of alleles, drawing on known repeat motifs from fact sheets. This step converts the allele sequences into bracket sequences, thereby simplifying the identification of incomplete units, SNPs, or InDels. A key advantage of NASTRA over alignment-based methods is its independence from the allele reference database, enabling it to genotype STRs even when allele information is missing from the database. Moreover, we used this method on MiSeq FGx data during our benchmark construction, proving its applicability to NGS data as well.

To comprehensively evaluate NASTRA’s performance, we conducted a series of assessments involving various amplification kits and flow cells. These include analyses of ForenSeq data, PowerSeq data, and two practical case studies. Initially, we created a benchmark dataset comprising 84 DNA samples using the MiSeq FGx system, a widely accepted technology for forensic applications. Then, we expanded the training and test sets to obtain the optimal thresholds for NASTRA. Further, we compared NASTRA with the alignment-based tools STRspy on ForenSeq data and PowerSeq data in terms of accuracy and speed. It’s important to note that we did not include renowned repeat quantification tools like repeatHMM (24), STRique (25), and DeepRepeat (26) in our comparison. These tools are specifically designed for detecting repeat expansion disorders and may not provide the accuracy needed for forensic STR genotyping. Additionally, we conducted two case studies: cell line STR profiling and paternity testing. In summary, our results demonstrate that NASTRA excels in accuracy and speed, underscoring its potential as an effective method for real-world applications.

However, NASTRA has several limitations for further improvement. Firstly, the comparison was limited to only one alignment-based method, and the associated reference database is still under development. Consequently, this comparison might not offer a completely objective assessment but rather a representative illustration in terms of accuracy and runtime. Secondly, although NASTRA can detect incomplete repeat units in the output allele, it does not integrate this digital information into its allele calls. Thirdly, there’s a risk in NASTRA’s read clustering process that could lead to two alleles being misclassified as one, particularly in cases involving a single nucleotide polymorphism (SNP) or a one-base gap between two similar alleles. Fourthly, our tests on ForenSeq and PowerSeq data revealed that amplicon length influences NASTRA’s performance. Moreover, the similarity in the flanking regions of STRs, including prefixes and suffixes, could result in incorrect trimming. Therefore, for accurate STR genotyping, NASTRA requires amplicons with extended flanking regions (over 30 bp). Lastly, NASTRA’s effectiveness is contingent on the quality of base sequences input, as it relies heavily on the base calling model. Given that STRs are complex genomic regions, developing a robust basecalling model specifically for STRs is imperative.

NASTRA represents an innovative computational approach for STR genotyping in nanopore sequencing data, employing a structure-aware algorithm that bypasses traditional alignment with a reference database. Our evaluations, conducted using various flow cells and amplification kits, have demonstrated NASTRA’s proficiency in both genotyping accuracy and speed. While not all STR loci consistently produced correct genotype calls, NASTRA showed notable robustness in genotyping specific loci, underscoring its substantial potential for individual identification and cell line authentication in nanopore sequencing contexts. To further enhance NASTRA’s capabilities, additional validation studies with larger sample sizes are essential.

## DATA AVAILABILITY

NanoTime is available on GitHub (https://github.com/renzilin/NanoTime). NASTRA is under GPL v3.0 license and is publicly available on GitHub (https://github.com/renzilin/NASTRA). The data that support the findings of this study have been deposited into CNGB Sequence Archive (CNSA) (34) of China National GeneBank DataBase (CNGBdb) (35) with accession number CNP0004956.

## SUPPLEMENTARY DATA

Supplementary Data are available at NAR online.

## AUTHOR CONTRIBUTIONS

Zilin Ren: Conceptualization, Methodology, Data curation, Formal Analysis, Writing – original draft. Jiarong Zhang: Investigation, Validation. Tingting Yang: Investigation, Validation. Yixiang Zhang: Data curation, Software, Writing – review & editing. Pingping Sun: Writing – review & editing. Jiguo Xue: Writing – review & editing. Bo Zhou: Writing – review & editing. Jiangwei Yan: Resources, Supervision. Ming Ni: Supervision, Writing – review & editing.

## ACKNOWLEDGEMENTS

The authors would like to thank Dr. JinDing Liu, Dr. Fenglong Yang, and Dr. Juan Jia from Shanxi Medical University, as well as Dr. Xu Liu, for their valuable comments and support throughout this study.

