## supplementart materials for "NASTRA: Accurate analysis of short tandem repeat markers by nanopore sequencing with repeat-structure-aware algorithm"

**Supplementary Materials**

Since NASTRA relies on the motifs from STRbase for inferring repeated structures, the motifs it uses don't account for potential SNPs. Therefore, we have implemented adaptive adjustments in NASTRA for certain loci to ensure accurate genotyping. These adjustments focus on identifying SNPs in both the flanking regions and the repeat regions themselves. Below is a detailed description of these adjustments.

D13S317 is a simple TATC tetranucleotide repeat. In the 3′ flank adjacent to the repeat region is AATC AATC CTCA. Due to polymorphisms (1), the sequence can change to <u>T</u>ATC AATC CTCA or <u>T</u>ATC <u>T</u>ATC CTCA, potentially leading to misinterpretation of the TATC sequence as part of the repeat. To address this, NASTRA modifies its genotyping calls, taking into consideration the number of TATC sequences present in this adjacent area.

D5S818 is a simple ATCT tetranucleotide repeat. In the 5′ flank adjacent to the repeat region is CTCT. A SNP might alter the sequence from CTCT to ATCT (2), potentially leading to the misidentification of ATCT as part of the repeat. To accurately address this, NASTRA adjusts its genotyping calls by specifically identifying and accounting for the CTCT sequence.

D20S482 is a simple AGAT tetranucleotide repeat. In the 3′ flank close to the repeat region, the sequence contains AGCT. A possible SNP could change this to AGAT, which might then be mistakenly identified as part of the repeat sequence. To counter this issue, NASTRA modifies its genotyping calls by specifically recognizing and accounting for the AGCT sequence.

D2S1338 is characterized by a compound tetranucleotide repeat with a structure of [GGAA]n [GGAC]m [GGAA]x [GGCA]y. Polymorphisms can occur in [GGAA]n, leading to a transformation of GGAA into GAAA (3). To address this, NASTRA revises its genotyping calls by considering the length of nucleotides, including those not typically counted in the allele.

D2S441 contains a simple TCTA repeat. There are compound motifs consisting of a varying number of TCTA repeats, followed by a single tetranucleotide of either TCAA,

TTTA, or TCTG, and concluding with more TCTA repeats. The total count for these compound motifs varies from 10 to 17 repeats, in the format [TCTA]n [TNNN] [TCTA]n (3, 4). NASTRA modifies the genotyping calls by the length for the TNNN.
D4S2408 is characterized by a simple ATCT repeat, with the motif structured as [ATCT]n. However, due to the presence of SNPs within the motif, the repeat structure can vary to [ATCT]m GTCT [ATCT]n. In response to this variation, NASTRA adapts its genotyping calls by considering the length of nucleotides that are not typically included in the allele count.
D7S820 features a simple TCTA repeat with the motif represented as [TCTA]n. However, the presence of SNPs within this motif can alter the repeat structure to [TCTA]m CCTA [TCTA]n (5). In response to these variations, NASTRA adjusts its genotyping calls by considering the length of nucleotides that are typically not included in the allele count.
D18S51 is characterized by a simple AGAA repeat, with the motif typically structured as [AGAA]n. However, the presence of SNPs within this motif can lead to a variation in the repeat sequence, resulting in a structure like [AGAA]m AGCA [AGAA]n. To address this, NASTRA revises its genotyping calls, taking into account the length of nucleotides that are not conventionally included in the allele count.
D9S1122 features a simple TAGA repeat. The typical motif structure is [AGAA]n, but we have identified SNPs within this motif that can alter the repeat to [TAGA]1 TCGA [TAGA]n. Consequently, NASTRA modifies its genotyping calls by factoring in the length of nucleotides that aren't typically counted in the allele.
The vWA locus is known for its compound tetranucleotide repeat, with a possible repeat structure of [TAGA]x [CAGA]y [TAGA]z. Polymorphisms within [TAGA]n can lead to a conversion of TAGA into TGGA (6). In response to this, NASTRA adapts its genotyping calls by considering the length of nucleotides that are typically not included in the allele count.

1. Wang,L., Zhao,X.-C., Ye,J., Liu,J.-J., Chen,T., Bai,X., Zhang,J., Ou,Y., Hu,L., Jiang,B.-W., *et al.* (2014) Construction of a library of cloned short tandem repeat (STR)

alleles as universal templates for allelic ladder preparation. *Forensic Science International: Genetics*, **12**, 136–143.

### Supplementary Tables

**Supplementary Table S1.** The accuracy comparison between NASTRA and STRspy on ForenSeq data.

| Locus | NASTRA | STRspy |
|---|---|---|
| CSF1PO | 1 | 1 |
| D10S1248 | 1 | 1 |
| D12S391 | 1 | 1 |
| D16S539 | 1 | 1 |
| D2S441 | 1 | 1 |
| D3S1358 | 1 | 1 |
| D8S1179 | 1 | 1 |
| TH01 | 1 | 1 |
| TPOX | 1 | 1 |
| D5S818 | 1 | 0.973 |
| D7S820 | 1 | 0.938 |
| PentaE | 1 | 0.926 |
| D19S433 | 1 | 0.923 |

| | | |
|---|---|---|
| D21S11 | 1 | 0.921 |
| D13S317 | 1 | 0.718 |
| D22S1045 | 1 | 0.7 |
| vWA | 1 | 0.684 |
| D17S1301 | 1 | - |
| D20S482 | 1 | - |
| D4S2408 | 1 | - |
| D9S1122 | 1 | - |
| D1S1656 | 0.971 | 0.974 |
| D6S1043 | 0.971 | - |
| D2S1338 | 0.925 | 1 |
| PentaD | 0.875 | 1 |
| D18S51 | 0.8 | 0.825 |
| FGA | 0.784 | 0.816 |

**Supplementary Table S2.** The accuracy comparison between NASTRA and STRspy on PowerSeq data.

| Locus | NASTRA | STRspy |
|---|---|---|
| CSF1PO | 1 | 1 |
| D10S1248 | 1 | 1 |
| D12S391 | 1 | 1 |
| D22S1045 | 1 | 1 |
| D2S441 | 1 | 1 |
| D3S1358 | 1 | 1 |
| D5S818 | 1 | 1 |
| D8S1179 | 1 | 1 |
| TPOX | 1 | 1 |
| D16S539 | 1 | 0.978 |
| D19S433 | 1 | 0.978 |
| D1S1656 | 1 | 0.978 |
| D2S1338 | 1 | 0.956 |
| D21S11 | 1 | 0.933 |
| D7S820 | 1 | 0.927 |
| D13S317 | 1 | 0.783 |
| vWA | 1 | 0.63 |
| TH01 | 0.978 | 1 |
| FGA | 0.935 | 0.891 |
| PentaE | 0.914 | 0.919 |
| D18S51 | 0.911 | 0.667 |
| PentaD | 0.833 | 1 |

**Supplementary Table S3.** Throughput per sample across different sequencing durations in the paternity test. Notably, the throughput values are calculated using the entire sequencing data from the ForenSeq amplicons, even though some parts of this data are not utilized by NASTRA.

| | | Samples | | | | | |
|---|---|---|---|---|---|---|---|
| | **throughput (Mbp)** | 2800M | 9948 | C | F | M | GM |
| **Sequencing durations** | 6 min | 1.90 | 2.52 | 2.35 | 2.74 | 1.84 | 1.96 |
| | 12 min | 4.27 | 5.59 | 5.14 | 6.12 | 4.02 | 4.43 |
| | 18 min | 6.58 | 8.59 | 7.89 | 9.42 | 6.16 | 6.89 |
| | 24 min | 8.71 | 11.56 | 10.53 | 12.65 | 8.16 | 9.28 |
| | 30 min | 10.80 | 14.38 | 13.04 | 15.78 | 10.08 | 11.48 |
| | 1h | 20.64 | 26.95 | 24.41 | 29.25 | 18.38 | 21.26 |
| | 2h | 34.83 | 46.76 | 43.43 | 50.26 | 31.15 | 36.48 |
| | 3h | 47.64 | 64.69 | 59.44 | 69.42 | 42.50 | 50.38 |
| | 4h | 58.70 | 80.04 | 73.01 | 85.84 | 52.29 | 62.18 |
| | 5h | 64.94 | 88.35 | 81.54 | 95.50 | 58.48 | 69.67 |

### Supplementary Figures

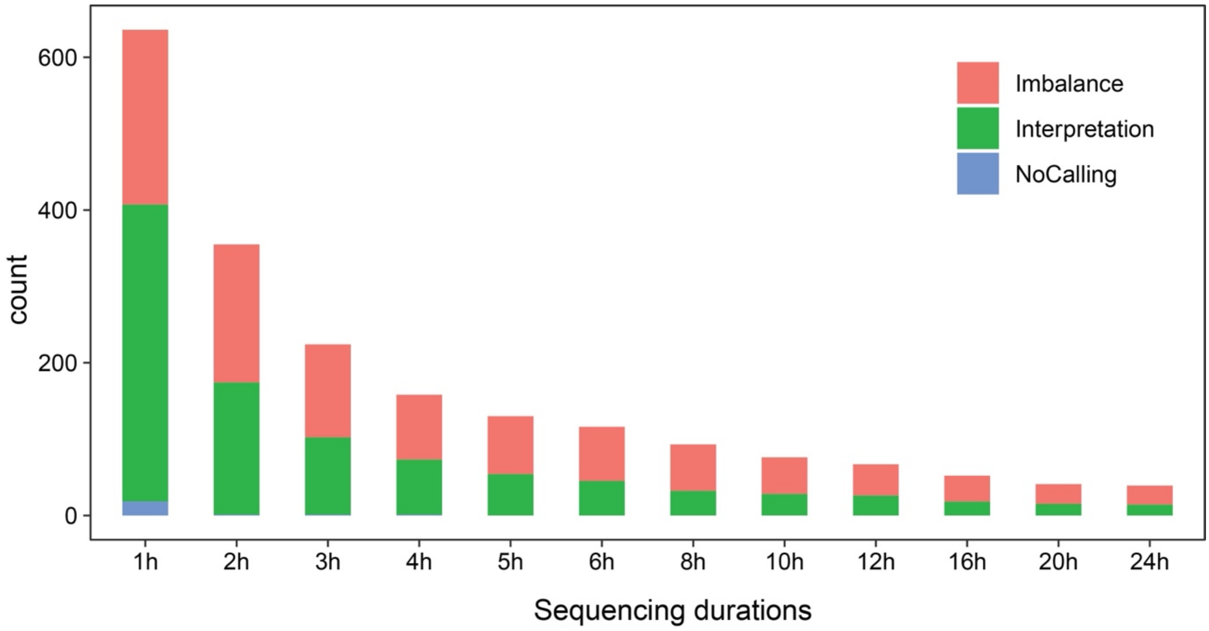


**Supplementary Figure S1.** The number of failed genotyping results with various sequencing duraitons in downsampled data. Blue represents for NoCalling, which indicates that the number of reads spanning the corresponding region is fewer than

10. Red represents for Imbalance, indicating the SN of minor allele is less than 25. Green represents for Interpretation, indicating the SN of major allele is less than 25.

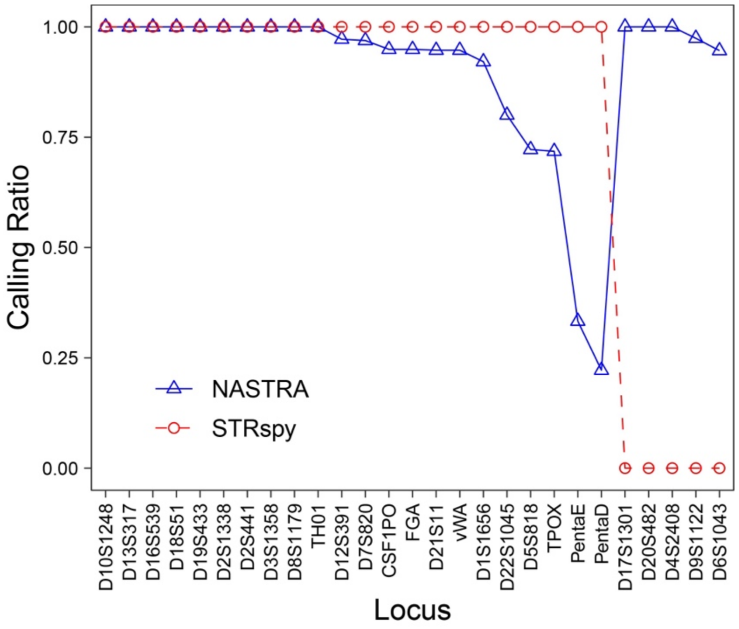


**Supplementary Figure S2.** The calling ratio of NASTRA and STRspy in ForenSeq data. For STRspy, the allele information for D17S1301, D20S482, D4S2408, D9S1122, and D6S1043 is absent from the provided allele reference.

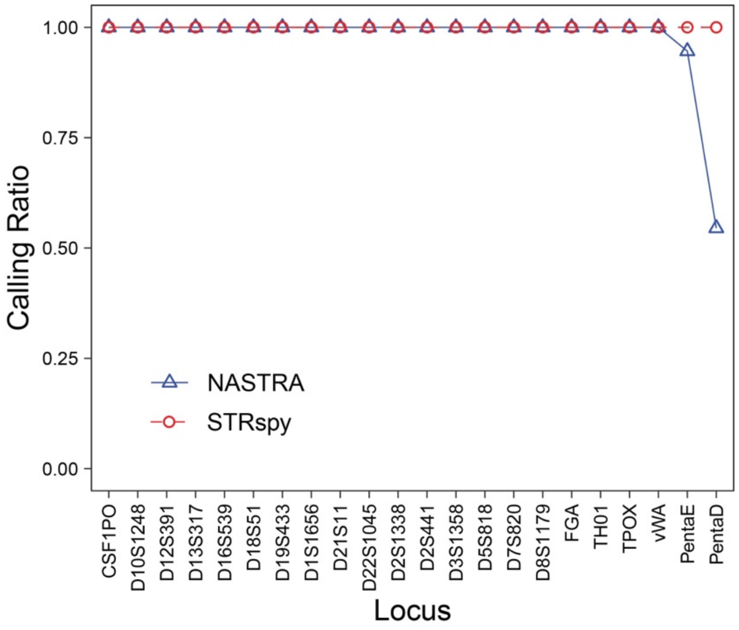


**Supplementary Figure S3.** The calling ratio of NASTRA and STRspy in PowerSeq data.

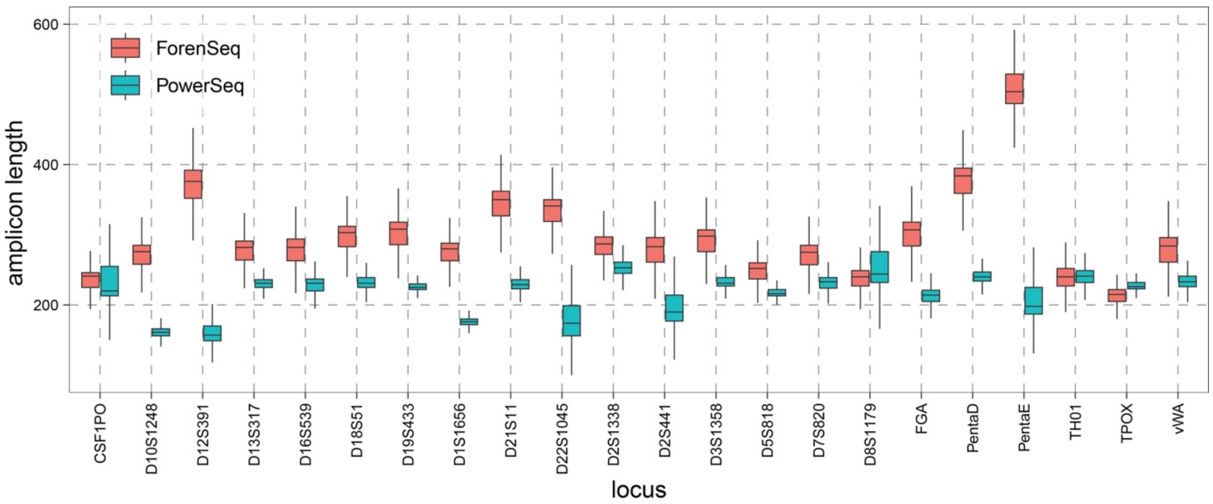


**Supplementary Figure S4.** The length distribution of amplicons in (a) ForenSeq data and (b) PowerSeq data. Overall, the amplicon length of ForenSeq data is slightly longer than that of PowerSeq data.
